# Nanobody-dependent delocalization of endocytic machinery in Arabidopsis root cells dampens their internalization capacity

**DOI:** 10.1101/2020.02.27.968446

**Authors:** Joanna Winkler, Andreas De Meyer, Evelien Mylle, Peter Grones, Veronique Storme, Daniël Van Damme

## Abstract

Plant cells perceive and adapt to an ever-changing environment by modifying their plasma membrane (PM) proteome. Whereas secretion deposits new integral membrane proteins, internalization by endocytosis removes membrane proteins and associated ligands, largely with the aid of adaptor protein complexes and the scaffolding molecule clathrin. Two adaptor protein complexes function in clathrin-mediated endocytosis at the PM in plant cells, the heterotetrameric Adaptor Protein 2 (AP-2) complex and the octameric TPLATE complex (TPC). Whereas single subunit mutants in AP-2 develop into viable plants, genetic mutation of a single TPC subunit causes fully penetrant male sterility and silencing single subunits leads to seedling lethality. To address TPC function in somatic root cells, while minimizing indirect effects on plant growth, we employed nanobody-dependent delocalization of a functional, GFP-tagged TPC subunit, TML, in its respective homozygous genetic mutant background. In order to decrease the amount of functional TPC at the PM, we targeted our nanobody construct to the mitochondria and fused it to TagBFP2 to visualize it independently of its bait. We furthermore limited the effect of our delocalization to those tissues that are easily accessible for live-cell imaging by expressing it from the PIN2 promotor, which is active in root epidermal and cortex cells. With this approach, we successfully delocalized TML from the PM. Moreover, we also show co-recruitment of TML-GFP and AP2A1-TagRFP to the mitochondria, suggesting that our approach delocalized complexes, rather than individual adaptor complex subunits. In line with the specific expression domain, we only observed minor effects on root growth, yet realized a clear reduction of endocytic flux in epidermal root cells. Nanobody-dependent delocalization in plants, here exemplified using a TPC subunit, has the potential to be widely applicable to achieve specific loss-of-function analysis of otherwise lethal mutants.

## 1 Introduction

Cells are delineated by their plasma membrane (PM). The PM houses a plethora of proteins ranging from receptors and ion channels to structural membrane proteins. Many of these PM proteins, commonly termed cargo, are responsible for cellular communication with the outside world. In eukaryotes, endocytosis is the cellular process where cargoes, associated ligands as well as lipids are internalized from the PM. Endocytosis thereby provides a way to regulate the content and consequently modulate protein activity at the PM. A predominant and well-studied form of endocytosis is clathrin-mediated endocytosis (CME) (Bitsikas et al., 2014). CME refers to the dependency of the scaffolding protein clathrin, which coats the developing and mature vesicles (Robinson, 2015). In plants, CME plays a role in hormone signaling (Irani et al., 2012; Martins et al., 2015; Zhang et al., 2017), nutrient availability (Wang et al., 2017; Dubeaux et al., 2018; Yoshinari et al., 2019), pathogen defense and susceptibility (Mbengue et al., 2016; Li and Pan, 2017), and other biotic and abiotic stresses (Li et al., 2011). Consequently, CME is essential for plant development.

Two early-arriving adaptor complexes, the heterotetrameric Adaptor Protein-2 complex (AP-2) and the hetero-octameric TPLATE complex (TPC) facilitate CME in plants. In contrast to AP-2, TPC represents an evolutionary ancient protein complex, which is lost in yeast and mammalian cells (Hirst et al., 2014). The slime mold *Dictyostelium discoideum* contains a similar complex, named TSET. TSET however is a hexameric complex in contrast to TPC in *A. thaliana*, which has two additional subunits. Also contrary to TPC, TSET is dispensable in *D. discoideum* (Hirst et al., 2014). The presence of a full or partial TSET complex in other eukaryotes was confirmed by additional homology searches, indicative of its ancient evolutionary origin (Hirst et al., 2014).

AP-2 and TPC have both common and distinct functions, possibly relating to cargo specificity and/or fate of the internalized cargo (Bashline et al., 2015; Sánchez-Rodríguez et al., 2018; Wang et al., 2019; Yoshinari et al., 2019). In addition, functional diversification of both complexes is reflected in their mutant phenotypes. Knockout plants in individual AP-2 subunits are affected at various stages of development but viable (Di Rubbo et al, 2013; Kim et al, 2013; Fan et al, 2013; Yamaoka et al, 2013; Bashline et al, 2013). However, *ap2* mutants show reduced internalization of the styryl dye FM4-64, which can be seen as proxy to a difference in cargo uptake (Jelínková et al., 2010), as well as known endocytic cargoes like the brassinosteroid receptor BRASSINOSTEROID INSENSITIVE 1 (BRI1), the Boron exporter BOR1 and auxin efflux carriers of the PIN family (Di Rubbo et al., 2013; Fan et al., 2013; Kim et al., 2013; Yoshinari et al., 2016, 2019).

The relatively mild phenotype of *ap2* single subunit mutants in plants contrasts with the lethal phenotype of a single *ap2* subunit knockout in mice (Mitsunari et al., 2005). Alternatively, the complex does not seem to be essential for yeast (Yeung et al., 2013). In *Caenorhabditis elegans*, AP-2 subunits are capable of assembling into hemicomplexes which partially retain their functionality (Gu et al., 2013). In plants, AP2M and AP2S are still recruited to the PM in *ap2s* and *ap2m* mutants respectively (Wang et al., 2016), suggesting that AP-2 hemicomplexes might also confer partial functionality in plants.

In contrast to AP-2, single knockouts of TPC subunits result in fully penetrant male sterility with shriveled pollen and ectopic callose accumulation (Van Damme et al., 2006; Gadeyne *et al*, 2014). Similar pollen-lethal phenotypes are also reported for *drp1c* (Backues et al., 2010) as well as *clc1* (Wang et al., 2013), involved in vesicle fission and clathrin triskelion assembly respectively.

So far, there is only one viable weak allele of one TPC subunit identified. This *twd40-2-3* mutant (Bashline et al., 2015) is however likely merely a knockdown as *twd40-2-1* and *twd40-2-2* mutants are pollen lethal (Gadeyne et al., 2014). Knockdowns of *TML* and *TPLATE* resulted in seedling lethality with a reduced internalization of FM4-64, BRI1, RECEPTOR-LIKE PROTEIN 44 (RLP44) and the cellulose synthase subunit CESA6 (Irani et al., 2012; Gadeyne et al., 2014; Sánchez-Rodríguez et al., 2018; Gómez et al., 2019). Silencing works on the messenger level and phenotypes only become apparent following degradation of pre-made proteins. As adaptor protein complexes can be recycled following each round of internalization, approaches affecting these complexes at the protein level have a more direct effect. In animal cells, conditional delocalization using rapamycin to target AP-2 to mitochondria has been successfully applied to interfere with endocytosis (Robinson et al., 2010).

Since their discovery, nanobodies, derived from camelid heavy chain-only antibodies (HCAb), have found their way into a wide variety of applications in biological fields. Nanobodies are similar to antibodies (Ab) in the sense that they can bind epitopes with high affinity in a highly selective manner (Ingram et al., 2018). Their applications range from drug discovery, crystallography and imaging techniques to probing protein functions (Ingram et al., 2018). The latter can be done by enforcing nanobody-dependent protein degradation or nanobody-dependent localization (Caussinus et al., 2012; Früholz et al., 2018; Ingram et al., 2018). Nanobodies can be expressed as a single chain, compact and stable protein while still retaining high selectivity and affinity for its epitope (Muyldermans, 2013). This makes them more convenient to clone and to express compared to conventional antibodies.

A nanobody-dependent method, degradFP, was developed in *Drosophila melanogaster*, to generate a conditional knockout at the protein level. This tool uses an anti-GFP nanobody, linked to an F-box to target it for ubiquitin-dependent degradation (Caussinus et al., 2012). This approach has also very recently been successfully used in plants to degrade WUSCHEL-GFP (Ma et al., 2019). Nanobodies have also been used in Arabidopsis seedlings to lock down vacuolar sorting receptors (VSRs) in cellular compartments upstream of TGN/EE, allowing to determine their retrograde trafficking pathway (Früholz et al., 2018).

Finally, nanobody-dependent lockdown was successfully applied in HeLa cells where EPS15, a pioneer endocytic accessory protein (EAP) that facilitates initiation of CME by stabilizing AP-2 presence at the PM, was successfully delocalized by expressing an anti-EPS15 nanobody on endosomes or mitochondria, thereby inactivating it (Traub, 2019).

Lock down of proteins to a cellular compartment of choice and can thus be effectively used in a similar fashion as the rapamycin-based system from the Robinson lab (Robinson et al., 2010). Here, we explore the effects on CME by delocalizing a GFP-tagged functional TML-GFP fusion protein to the mitochondria in the homozygous *tml-1(-/-)* mutant background using a nanobody directed against eGFP.

## 2 Results

### 2.1 A mitochondrially targeted nanobody can delocalize TML

TPC is a robust multi-subunit complex functioning at the PM and TPC can be affinity purified using any of its subunits as bait (Gadeyne et al., 2014). In order to delocalize, and thereby inactivate TPC, we took advantage of the functionally complemented homozygous *tml-1*(-/-) mutant expressing TMLprom::TML-GFP (Gadeyne et al., 2014). In this background, we introduced expression of a nanobody directed against eGFP (GFPNb) (Künzl et al., 2016), which we visualized by fusing it to TagBFP2. We targeted the fusion protein to the mitochondria using the import signal of the yeast mitochondrial outer membrane protein Tom70p as described before (Robinson et al., 2010). This targeting signal is functional in plants as we have previously colocalized constructs containing this signal with mitoTracker in *N. benthamiana* leaf epidermal cells (Winkler et al., unpublished results). We used the PIN2prom to drive expression of MITOTagBFP2-GFPNb in epidermis and cortex root files, which are easy to image with respect to future experiments. MITOTagBFP2-GFPNb localized to discrete punctae in Arabidopsis wild type roots (**Figure 1A**). These punctae appeared to have different sizes, with the large ones likely representing clusters. Co-staining with the mitochondrial dye MitoTracker Red revealed hardly any colocalization (**Figure 1A**), which might suggest that expression of MITOTagBFP2-GFPNb has an effect on mitochondrial fitness. Nevertheless, we used this tool to attempt to delocalize TML away from the PM.

**Figure 1.**
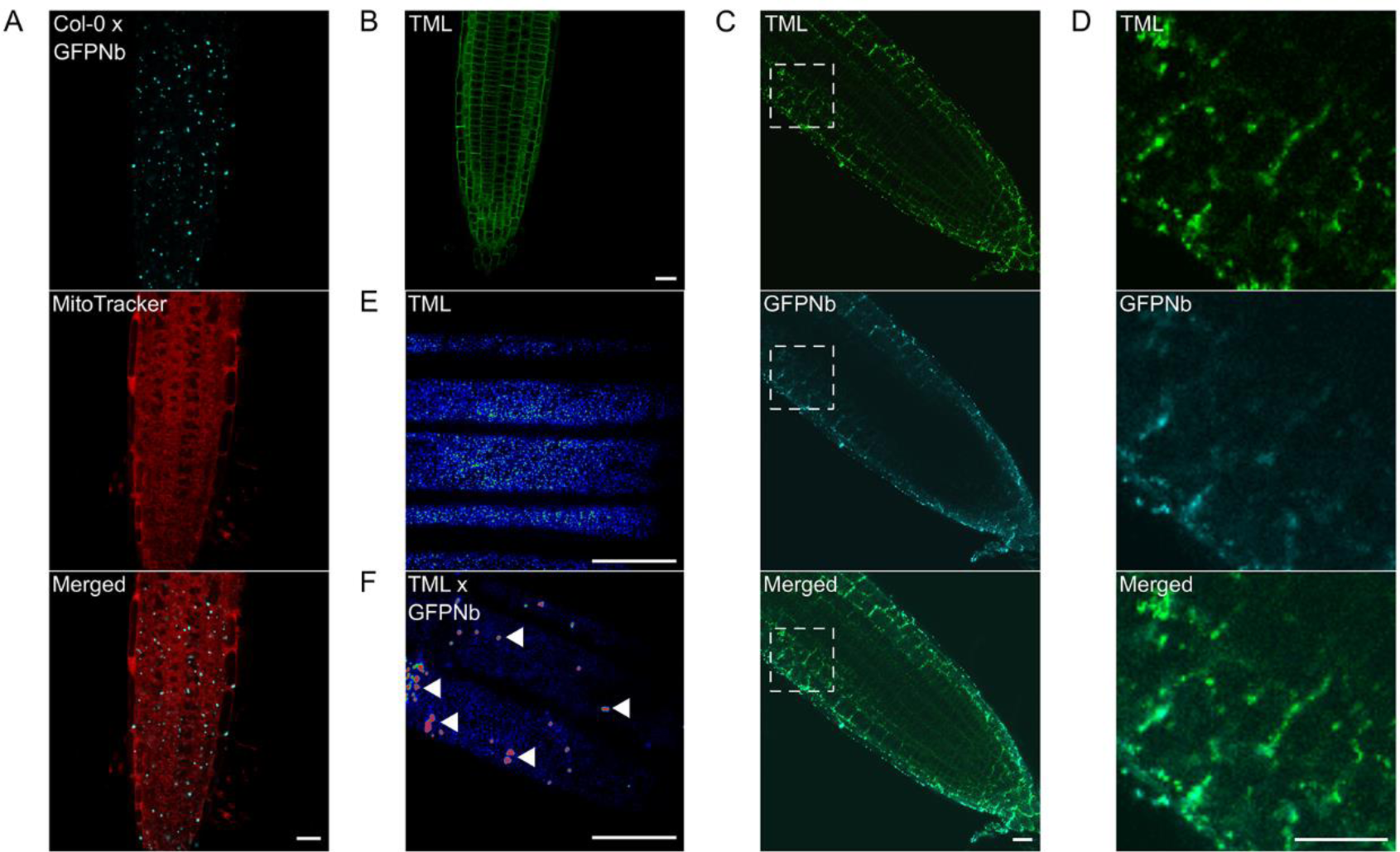
Expression of a mitochondrial-targeted nanobody against GFP allows delocalization of TML-GFP. (A) Representative image of a wild type root expressing MITOTagBFP2-GFPNb counterstained with MitoTracker Red showing targeting of the construct to cytosolic punctae of various sizes, likely representing dysfunctional clustered mitochondria. (B) Representative Arabidopsis root image of *tml-1(-/-)* complemented with TML-GFP showing that the functional TML fusion is predominantly targeted to the PM. (C and D) Representative overview images and respective blow-ups of the outlined region of Arabidopsis roots where TML-GFP in *tml-1(-/-)* was combined with MITOTagBFP2-GFPNb expression, leading to its delocalization from the PM. (E and F) Representative, rainbow intensity colored, grazing sections through the PM, showing the recruitment of TML to endocytic foci without (E) and with partial delocalization of TML-GFP (F, arrowheads). Scalebars equal 20 µm.

In complemented *tml-1*(-/-) Arabidopsis roots, TML-GFP is recruited predominantly at the plasma membrane in a single confocal section (**Figure 1B**). Combining this line with the GFPNb, whose expression was restricted to the root epidermis and cortex files (**Figure 1C**), led to a change in the uniform plasma membrane labeling of TML to a denser staining of discrete punctae in these cell files. Most of those were still near the plasma membrane and colocalized with the fluorescent signal from the nanobody, indicating effective delocalization of TML-GFP (**Figure 1C and enhanced in 1D**). This delocalization was not apparent in the deeper layers of the root, where TML remained uniformly recruited to the plasma membrane (**Figure 1C**). Detailed analysis using spinning disk microscopy confirmed the strong recruitment of TML to those mitochondria that were present in the focal plane of the PM (**Figure 1E and 1F, arrowheads**). Next to the mitochondria however, TML was still recruited to endocytic foci at the plasma membrane in root epidermal cells. The density of endocytic foci in epidermal root cells is very high (Dejonghe et al., 2016, 2019; Sánchez-Rodríguez et al., 2018) and the density of endocytic foci, marked by TML-GFP, appeared similar between epidermal cells in the complemented mutant (control) background and in those cells that in addition also expressed GFPNb. The fluorescence intensity of the foci was however markedly reduced, in agreement with a substantial amount of TML-GFP accumulating at the mitochondria (**compare Figure 1E and 1F**).

### 2.2 Nanobody-dependent delocalization of TML also affects other endocytic players

In plants, the heterotetrameric AP-2 complex and the octameric TPLATE complex are presumed to function largely, but not exclusively, together to execute CME (Gadeyne et al., 2014; Bashline et al., 2015; Wang et al., 2016; Adamowski et al., 2018). Both TPC and AP-2 have been shown to be involved in the internalization of cellulose synthase (CESA) complexes or the Brassinosteriod receptor BRI1 for example (Bashline et al., 2013, 2015; Di Rubbo et al., 2013; Gadeyne et al., 2014; Sánchez-Rodríguez et al., 2018).

Moreover, a joint function is also suggested from proteomics analyses, which could identify subunits of both complexes when the AtEH1/Pan1 TPC subunit was used as bait in tandem-affinity purification assays (Gadeyne et al., 2014). To investigate whether our tool, aimed at delocalizing TPC, would also interfere with AP-2 recruitment at the PM, we tested the localization of AP-2 when TML was targeted to the mitochondria. To do so, we crossed our TML-GFP line, *in tml-1*(-/-) and expressing PIN2prom::MITOTagBFP2-GFPNb with the homozygous complemented *tml-1*(-/-) line, expressing TML-GFP as well as one of the large AP-2 subunits, AP2A1, fused to TagRFP (Gadeyne et al., 2014). Offspring plants that did not inherit the nanobody construct showed PM and cell plate recruitment of TML and AP2A1, and only background fluorescence in the TagBFP2 channel (**Figure 2A**). In the offspring plants that inherited the nanobody construct however, the localization of the adaptor complex subunits changed. Both TML and AP2A1 accumulated at punctae, which clearly colocalized with the TagBFP2-fused nanobody construct (**Figure 2B**). The observed delocalization of AP2A1 to the mitochondria, together with TML strongly suggests that our approach has the capacity to delocalize TPC and AP-2 rather than TML alone, given that TPC and AP-2 are presumed to be linked via the AtEH1/Pan1 subunit (Gadeyne et al., 2014).

**Figure 2.**
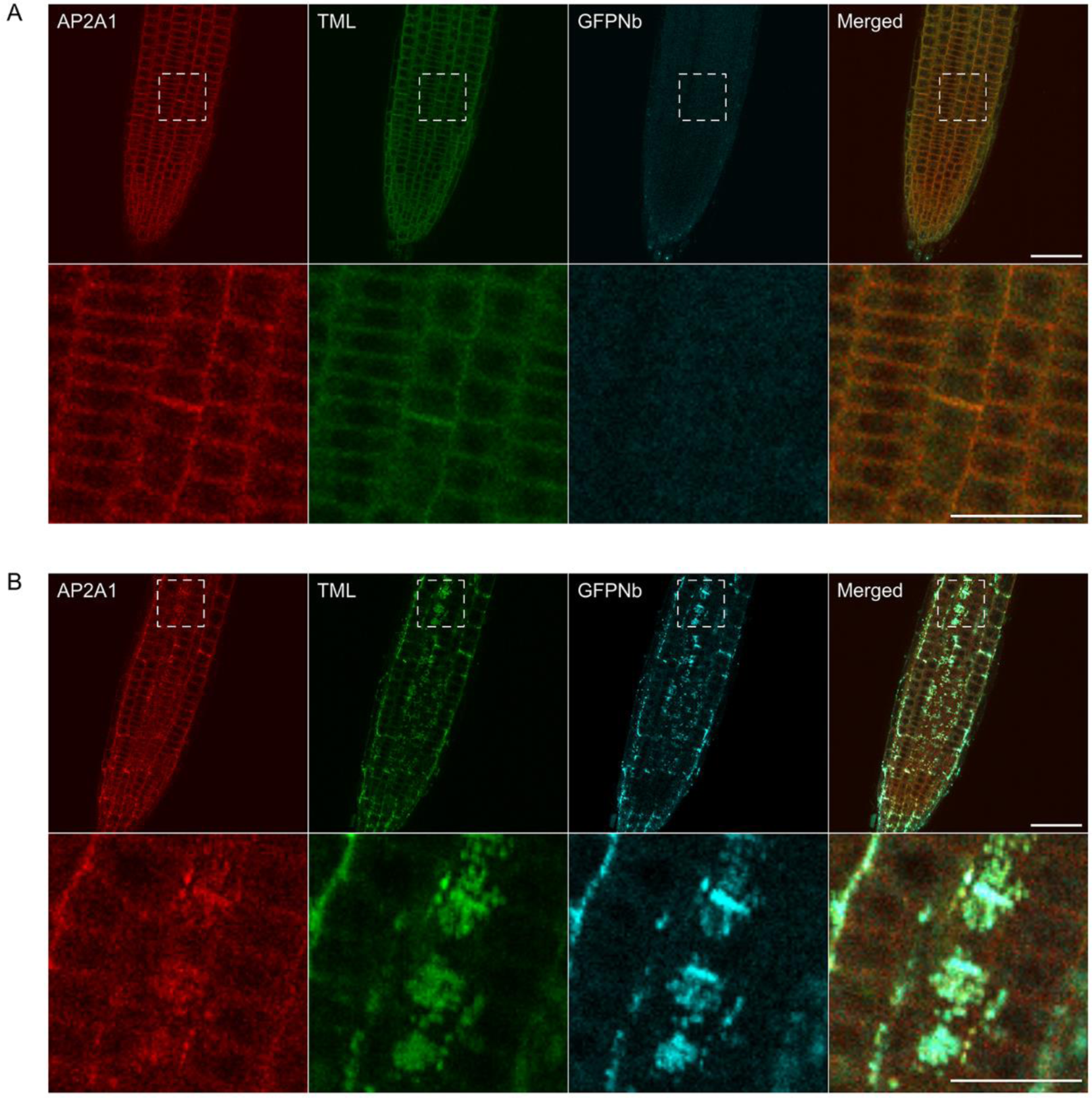
Delocalization of TML also affects the targeting of other endocytic players. (A and B) Representative images and blow-ups of the outlined regions of Arabidopsis roots expressing TML-GFP and AP2A1-TagRFP without (A) and with (B) MITOTagBFP2-GFPNb expression. GFPNb expression causes delocalization of both TML and AP2A1. Scale bars equal 20 µm (overview pictures) or 10 µm (blow up pictures).

### 2.3 Mistargeting adaptor complexes in epidermis and cortex affects root endocytic uptake with only minor effects on root growth

In contrast to AP-2, genetic interference with TPC subunits causes fully penetrant male sterility (Van Damme et al., 2006; Di Rubbo et al., 2013; Fan et al., 2013; Kim et al., 2013; Yamaoka et al., 2013; Gadeyne et al., 2014). TPC functionality therefore requires all subunits, and constitutive homozygous loss-of-function backgrounds are therefore non-existing. Abolishing endocytosis in plants, by silencing TPC subunits (Gadeyne et al., 2014) or over expression of the uncaging proteins AUXILLIN-LIKE 1 or 2 (Adamowski et al., 2018) severely affects seedling development. The effect of silencing TPC subunits only indirectly affects protein levels and targeting clathrin might interfere with trafficking at endosomes besides the PM. As TPC and AP-2 only function at the PM, inactivating their function should not directly interfere with more downstream aspects of endosomal trafficking. Furthermore, by restricting the expression domain where adaptor complex function is tuned down to the two outermost layers in the root should allow to study internalization from the PM, independently of possible indirect effects caused by the severe developmental alterations.

We evaluated the growth of several different lines expressing either GFPNb alone: Col-Nb1 (*+/-*) and Col-Nb2 (*-/-*), or GFPNb combined with TML-GFP in the complemented *tml-1(-/-)* mutant background: TML-Nb1 (*-/-*) and TML-Nb2 (*-/-*), however no major developmental defects were observed (**Figure 3A**). Root length measurements of light grown seedlings revealed growth enhancement in lines Col-Nb2 compared to Col-0 and TML-Nb2 compared to TML (**Figure 3B**). Variability in between the different independent lines is most probably the result of GFPNb expression levels. These results suggest that nanobody expression and partial delocalization of TML have no negative effect on seedlings development under normal growth conditions.

**Figure 3.**
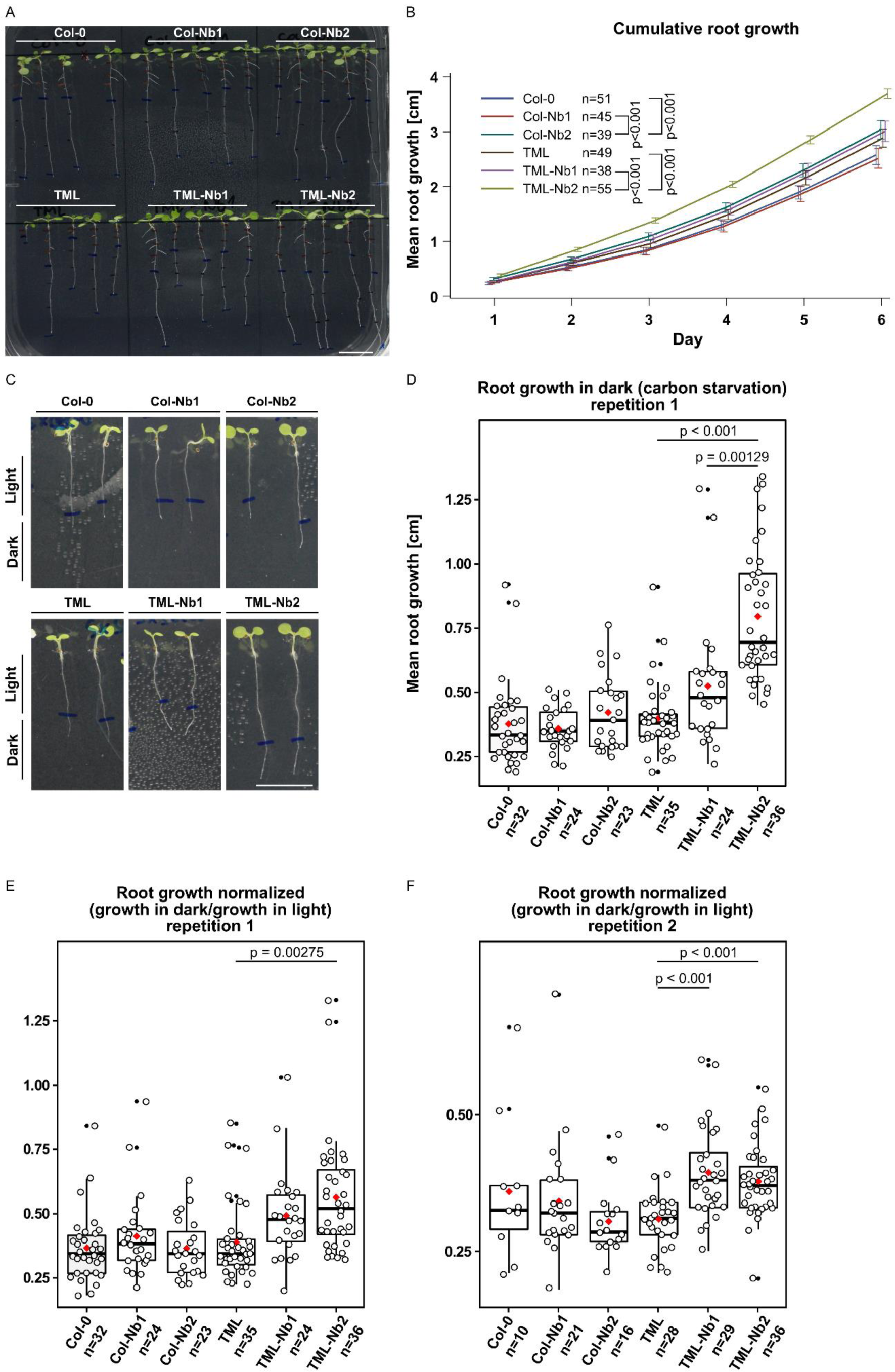
Delocalizing TML-GFP in root epidermal and cortical cells has only minor effects on root growth. Comparison of wild type seedlings (Col-0), wild type seedlings expressing MITOTagBFP2-GFPNb (Col-Nb1 and Col-0-Nb2) complemented *tml-1(-/-)* mutants expressing TML-GFP (TML) and complemented *tml-1(-/-)* mutants expressing TML-GFP and MITOTagBFP2-GFPNb (TML-Nb1 and TML-Nb2) in different light conditions. (A and B) Representative images of seedlings and quantification of root growth in continuous light. The quantification shows a cumulative root growth curve for each line and error bars represent the 95% confidence interval. p-values indicated were determined via mixed model statistics. (C-F) Representative images of seedlings grown for 5 days in continuous light and subsequently for 7 days in continuous dark. Measurements of growth in dark, as well as the respective dark/light ratio are represented in jitter box plot representations (the lines represent the median and the diamonds represent the mean). The statistical significance was determined using the Tukey contrasts procedure for Comparing Multiple Means under Heteroscedasticity. Number of individual seedlings analyzed per group is represented by n. Scale bars equal 1 cm.

The AtEH/Pan1 TPC subunits were recently implicated in growth under nutrient-depleted conditions as downregulation of *AtEH1/Pan1* expression rendered plants hyper-susceptible to carbon starvation (Wang et al., 2019). We therefore assessed if delocalizing TML-GFP, as well as other endocytic players, would also render these plants susceptibility to nutrient stress. To do so, we measured root lengths of seedlings grown for five days in continuous light and afterwards we placed them in the dark for an additional seven days. Measurements of root growth in dark under carbon stress conditions did not show any differences between WT and Col-Nb lines (**Figure 3C, D**). However, both TML-Nb lines exhibited increased root growth compared to TML (**Figure 3C, D)**. We calculated the ratio of root growth in dark over root growth in light to avoid overestimation of the results due to extraordinary growth of GFPNb lines. The ratios revealed that sequestering TML in TML-Nb lines does not cause any negative effect, but rather is beneficial towards the root growth under nutrient-depleted environment (**Figure 3E**). We repeated the experiment and obtained similar results also for the second time (**Figure 3F**). Overall, the effects of TML relocalization did not reveal any severe defects on seedlings development, even on nutrient-depleted media.

The subtle differences observed by comparing the effect of delocalization of TML on plant growth are likely a consequence of the restricted expression domain of GFPNb. We therefore monitored the effects of delocalizing TML more directly by visualizing the internalization of the styryl dye FM4-64, which in plants is commonly used as proxy for endocytic flux (Rigal et al., 2015; Jelínková et al., 2019). To rule out indirect effects of targeting GFPNb to the mitochondria, we compared endocytic flux between Col-Nb1, TML-GFP *in tml-1*(*-/-*),TML-Nb1 (*-/-*) and TML-Nb2 (*-/-*). We observed a slight decrease in endocytic flux when comparing wild type seedlings with the complemented *tml-1(-/-)* line and a strong reduction in endocytic flux between the complemented mutant and both complemented mutant lines where TML was partially delocalized (**Figure 4A, B**). Direct visualization of endocytic flux therefore allowed us to conclude that expression of the PIN2prom::MITOTagBFP2-GFPNb has the capacity to interfere with endocytosis in Arabidopsis root epidermal cells and that this tool certainly has the capacity to generate knockdown, and maybe even knockout lines at the protein level.

**Figure 4.**
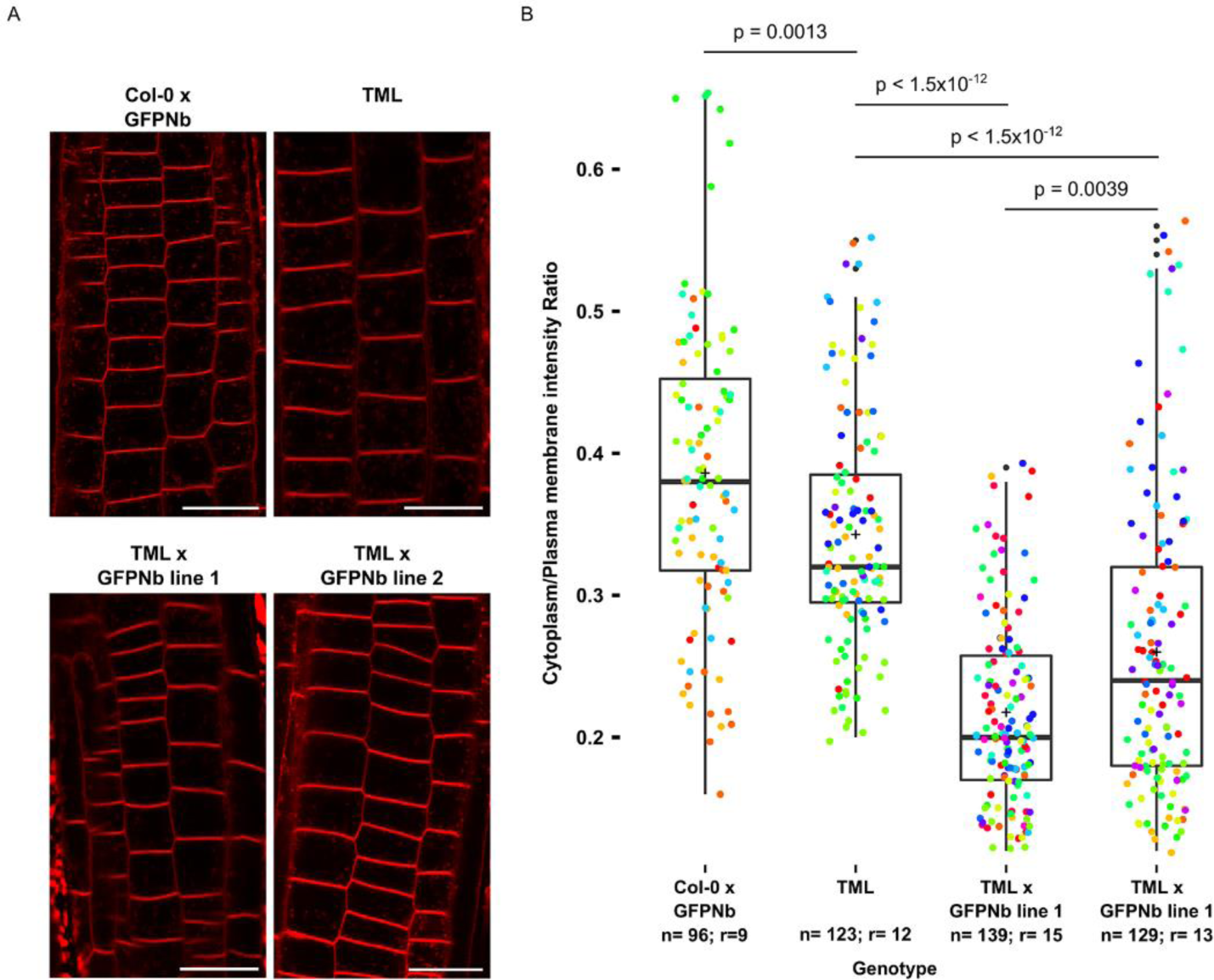
Nanobody-dependent delocalization reduces endocytic flux. (A) Representative single confocal slices of FM4-64 stained root cells of the different lines for which endocytic flux was quantified. FM4-64 uptake was compared between wild type Arabidopsis expressing MITOTagBFP2-GFPNb (Col-0 x GFPNb), the TML-GFP expressing complemented *tml-1(-/-)* mutant (TML), and two independent lines of the TML-GFP expressing complemented *tml-1(-/-)* mutant expressing MITOTagBFP2-GFPNb (TML x GFPNb). Scale bars equal 20 µm. (B) Box plot and Jitter box representation of the quantification of the cytoplasm/plasma membrane intensity of FM4-64 as proxy for endocytic flux. The black lines represent the median and the crosses represent the mean values. The dots represent individual measurements of cells. The rainbow-colored indication of the dots groups the cells from the different roots that were analyzed. The number of cells (n) and the number of individual roots (r) are indicated in the graph. The indicated p-values were calculated using the Wilcoxon-signed rank test.

## 3 Discussion

Analyzing how impaired TPC function directly affects endocytosis is hampered by the male sterility and/or seedling lethal mutant phenotypes following genetic interference of individual subunits (Gadeyne et al., 2014). Here, we explored to impair TPC function at the protein level by delocalizing a functional and essential subunit in its respective complemented mutant background. We were inspired by previous work in animal cells. However, instead of using rapamycin-dependent rerouting of one of the large AP-2 subunits, combined with silencing the endogenous subunit (Robinson et al., 2010), we took advantage of the complemented *tml-1(-/-)* mutant line expressing TML-GFP (Gadeyne et al., 2014) in combination with targeting a nanobody directed against GFP (GFPNb) (Künzl et al., 2016) to the mitochondria. We expressed the GFPNb in epidermis, cortex and lateral root cap as we expected ubiquitous constitutive expression to be lethal for the plant. Moreover, the epidermis and cortex cell files are easily accessible for imaging purposes. Proteins fused to this mitochondrial targeting signal colocalized with MitoTracker in transient *N. benthamiana* experiments (Winkler et al., unpublished results). This was not the case in Arabidopsis roots, indicating that constitutively decorating the mitochondria with the GFPNb construct affected their functionality without however causing a severe penalty on overall plant growth. The GFPNb system was capable of delocalizing TML-GFP and this caused the appearance of strongly fluorescent GFP-positive aggregations. Detailed inspection revealed however that our approach was insufficient to remove all TML from the PM. Compared to the control cells, sequestration of TML-GFP led to an overall reduction in signal intensity at the endocytic foci, without visually affecting their overall density. This also correlated with a significant reduction in endocytic tracer uptake, a proxy for reduced endocytosis. Intuitively, a reduced amount of complexes per endocytic spot would correlate with a weaker signal rather than a reduction in density. Our observation therefore fits with the occurrence and requirement of several TPC units to efficiently internalize a single clathrin coated vesicle.

The minor differences in root length, observed when TML-GFP was delocalized in the GFPNb lines, as well as the minor effects observed upon nutrient depletion growth can be explained by the limited expression domain of the PIN2 promoter. Increased root growth is nevertheless a mild effect compared to disrupting other parts of the CME machinery. Inducible overexpression of AUXILIN-LIKE1/2 results in complete seedling growth arrest with drastic effects on cell morphology (Adamowski et al., 2018). The same holds true for inducible expression of dominant-negative clathrin HUB and DRP1A (Kitakura et al., 2011; Yoshinari et al., 2016). Furthermore, estradiol-inducible TPLATE and TML knockdown lines are noticeably shorter and show bulging cells (Gadeyne et al., 2014). As we did not observe cellular effects in epidermal or cortical cell files, we conclude that our approach lacked the required strength to block endocytosis, but only reduced it.

Recent results suggest that plant cells very likely contain a feedback loop controlling TPC expression, as carbon starved plants contained roughly the same amount of full-length TPLATE-GFP, next to an extensive amount of TPLATE-GFP degradation products (Wang et al., 2019). In case plant cells make more TPC upon depleting the complex at the PM, DegradFP could provide a viable solution to this problem (Caussinus et al., 2012). By applying this method in GFP-complemented *tml-1(-/-)* mutants, newly synthesized TML-GFP would be broken down immediately, preventing to achieve functional levels of TPC at the PM. Stronger or inducible promotors and/or the use of a different targeting location might also increase the delocalization capacity. To avoid lethality due to ubiquitous sequestration, engineered anti-GFP nanobodies, whose affinity can be controlled by small molecules, could also be used (Farrants et al., 2020).

Untangling the function of TPC and AP-2 in CME at the plasma membrane requires tools that allow interfering specifically with the functionality of both complexes. Our nanobody-dependent approach targeting TPC via TML resulted in the co-delocalization of one of the large subunits of AP-2, indicating that we likely are not only targeting TPC, but also AP-2 function. Whether a complementary approach, by delocalizing AP-2, using AP2S or AP2M in their respective complemented mutant backgrounds, would also delocalize TPC is something that would be worth trying. Furthermore, as AP2S and AP2M subunits are still recruited in *ap2m* and *ap2s* single mutant backgrounds (Wang et al., 2016), AP-2 in plants might also function as hemi-complexes similar to what is reported in *C. elegans* (Gu et al., 2013). Single mutants therefore might not reflect functional null *ap2* mutants and a similar approach as performed here might also provide tools to inactivate AP-2 as a whole, which can be highly complementary to working with the single subunit mutants.

In conclusion, the data presented here is a first step toward the development of specific tools, which are required to help us understand the functions of AP-2 and TPC. On the long-term, this will generate insight into endocytosis at the mechanistic level and this will bring us closer to being able to modulate CME-dependent processes, and thereby modulating plant development, nutrient uptake as well as defense responses to our benefit.

## 4 Materials and Methods

### 4.1 Cloning

Gateway entry clones pDONR221-TagBFP2, pDONR221-MITOTagBFP2 and pDONRP2RP3-GFPNb were generated according to the manufacturer’s instructions (ThermoFisher Scientific BP clonase). pDONR221-TagBFP2 was amplified from pSN.5 mTagBFP2 (Pasin et al., 2014) with primers:

AttB1-GGGGACAAGTTTGTACAAAAAAGCAGGCTATGTCATCTAAGGGTGAAGAGCTTATCAAAGAGAAT and AttB2-GGGGACCACTTTGTACAAGAAAGCTGGGTCACCTCCGCCACCTCCACCTCCCAGTCCTGCGTA.

pDONR221-MITOTagBFP2 was generated from pDONR221-TagBFP2 by including the import signal of the yeast mitochondrial outer membrane protein Tom70p as described before (Robinson et al., 2010). The following primers sequences were used:

AttB1-GGGGACAAGTTTGTACAAAAAAGCAGGCTCAATGAAGAGCT TCATTACAAGGAACAAGACAGCCATTTTGGCAACCGTTGCTGCTACAGGTACTGCCATCGGTGCCTACTATTATTACAACCAATTGCAACAGGATCCACCGGTCGCCACCATGTCATCTAAGGGTGAAGAGCTT and AttB2-GGGGACCACTTTGTACAAGAAAGCTGGGTACGCTAAGTCTTCCTCTGAAATCAA.

pDONRP2RP3-GFPNb was generated from an anti-GFP Nanobody construct (Künzl et al., 2016) with primers attB2-GGGGACAGCTTTCTTGTACAAAGTGGGGATGTATCCTTATGATGTTC and attB3r-GGGGACAACTTTGTATAATAAAGTTGTTTAATGATGATGATGATGATGAGAAGA including a HA-tag, a 3xHis-tag and a stop codon.

The entry clones of the PIN2 promoter pDONRP4P1R_PIN2prom (Marquès-Bueno et al., 2016), pDONR221-MITOTagBFP2 and pDONRP2RP3-GFPNb were used in a triple Gateway LR reaction, combining pB7m34GW (Karimi et al., 2005) to yield pB7m34GW_PIN2prom::MITOTagBFP2-GFPNb.

### 4.2 Plant material and transformation

Plants expressing pB7m34GW_PIN2prom::MITOTagBFP2-GFPNb were generated by floral dip (Clough and Bent, 1998). Constructs were dipped into Col-0 and *tml-1(-/-)* (At5g57460) mutant lines described previously (Gadeyne et al., 2014). Primary transformants (T1) were selected on BASTA containing ½ strength MS medium without sucrose and 0.6% Gelrite (Duchefa, The Netherlands). PIN2prom::MITOTagBFP2-GFPNb expression was analyzed in the progeny of BASTA-resistant primary transformants (T2 seeds) by microscopy and T2 lines demonstrating strong expression were selected regardless of insert copy number. Next, T2 lines were crossed with the previously described TML-GFP complemented *tml-1(-/-)* mutant line expressing also RPS5Aprom::AP2A1-TagRFP (Gadeyne et al., 2014). Primary hybrids were analyzed via microscopy and best lines were selected on the basis of both PIN2prom::MITOTagBFP2-GFPNb and RPS5Aprom::AP2A1-TagRFP expression. For both Col-0 and *tml-1(-/-)* backgrounds, two independent lines (-Nb1 and -Nb2) were generated. Namely, Col-Nb1 (*+/-*), Col-Nb2 (*-/-*), TML-Nb1 (*-/-*) and TML-Nb2 (*-/-*).

### 4.3 Phenotypical quantification of root growth

Arabidopsis seedlings were grown at 21°C on ½ strength MS medium without sucrose and 0.6% Gelrite (Duchefa, Netherlands). For root growth plants were grown for 6 days in continuous light upon which the root growth of every seedling was marked. For carbon starvation, plants were grown for 5 days in continuous light after which the root growth of every seedling was marked Subsequently, the plates were covered and left for 7 days in dark after which root growth was marked again. Root growth and carbon starvation assays measurements were carried out with Fiji/ImageJ (Schindelin et al., 2012; Schneider et al., 2012). Statistical difference for root growth assay was determined via a mixed model analysis. Mixed linear model was applied to the root length of the lines Col-0, Col-Nb1, Col-Nb2, TML, TML-Nb1 and TML-Nb2 of each using the mixed procedure from SAS (SAS Studio 3.8 and SAS 9.4, SAS Institute Inc, Cary, NC). Fixed effects in the model were Line, Day and the interaction term. An unstructured covariance structure was estimated to model the correlations between measurements done on the same plant. The degrees of freedom of the fixed effects were approximated with the Kenward-Rogers method. The hypotheses of interests were the differences between Col-0 and its respectective nanobody lines, between TML and its respectective nanobody lines and between the nanobody lines with the same background. These hypotheses were tested using the plm procedure. p-values were adjusted for multiple testing using the maxT procedure as implemented in the plm procedure. Statistical difference for the carbon starvation assay was analyzed using Rstudio (Rstudio Team, 2019) with Welch corrected ANOVA to account for heteroscedasticity. Post hoc pairwise comparison was performed with the package MULTCOMP utilizing the Tukey contrasts (Herberich et al., 2010).

### 4.4 FM-uptake quantification

Endocytic tracer FM4-64 stock solution was prepared prior to treatment (2 mM in DMSO, Thermo Fisher). Roots were stained with 2 μM FM4-64 by incubating the seedlings in FM-containing ½ strength MS medium without sucrose for 30 min. Treatment was followed by microscopy. Acquired pictures were analyzed in Fiji/ImageJ (Schindelin et al., 2012; Schneider et al., 2012). PM and cytosol of individual epidermal cells were outlined (using the Select Brush Tool and Freehand selections, respectively) and histograms of pixel intensities were generated. Pictures which contained more than 1% saturated pixels were excluded from the quantification. Cytoplasm/PM ratios were calculated from average intensities of the top 1% highest intensity pixels based on the histograms. Outliers were removed via interquartile range in a single step. Data were analyzed using RStudio (Rstudio Team, 2019). Data distribution normality was check with Shapiro-Wilk test, and the significance level was tested with Wilcoxon-signed rank test for non-parametric data.

### 4.5 Image acquisition

Confocal images were taken using Leica SP8X confocal microscope equipped with a WLL laser and using the LASX software (Figure 1 A-D, Figure 2 and Figure 4). Images were acquired on Hybrid (HyD, gating 0.3-10.08 ns) and Photomultiplier (PMT) Detectors using bidirectional line-sequential imaging with a 40x water objective (NA=1.10) and frame or line signal averaging. Specific excitation and emission were used: 405nm laser and filter range 410-470nm for TagBFP2, 488nm laser and filter range 500-550nm for GFP, 488nm laser and filter range 600-740nm for FM4-64, 555nm laser and filter range 560-670 for TagRFP. Focal planes of plasma membranes (Figure 1E and 1F) were acquired with a PerkinElmer Ultraview spinning-disc system attached to a Nikon Ti inverted microscope and operated using the Volocity software package (Figure 1 E and F). Images were acquired on an ImagEMccd camera (Hamamatsu C9100-13) using frame-sequential imaging with a 100x oil immersion objective (NA=1.45). Specific excitation and emission was performed using a 488nm laser combined with a single band pass filter (500-550nm) for GFP and 405nm laser excitation combined with a single band pass filter (454-496nm) for TagBFP2. Images shown are single-slice.

## 6 Author Contributions

JW, ADM, EM and PG designed and performed experiments. DVD designed experiments and wrote the initial draft together with ADM. VS performed root growth assay statistical analysis. All authors contributed to the final version of the manuscript.

## 8 Acknowledgments

The authors would like to thank the ENPER members for forming a vibrant and open research community for more than 20 years already. We would also like to thank Steffen Vanneste (PSB, VIB/UGent, Belgium) for providing research tools. Research in the Van Damme lab is supported by the European Research Council (T-Rex project number 682436 to D.V.D., J.W. and A.D.M) and by the Research Foundation Flanders (FWO postdoctoral fellowship grant 1226420N to P.G.).

